# A crowdsourced analysis to identify *ab initio* molecular signatures predictive of susceptibility to viral infection

**DOI:** 10.1101/311696

**Authors:** Slim Fourati, Aarthi Talla, Mehrad Mahmoudian, Joshua G. Burkhart, Riku Klén, Ricardo Henao, Zafer Aydin, Ka Yee Yeung, Mehmet Eren Ahsen, Reem Almugbel, Samad Jahandideh, Xiao Liang, Torbjörn E.M. Nordling, Motoki Shiga, Ana Stanescu, Robert Vogel, The Respiratory Viral DREAM Challenge Consortium, Gaurav Pandey, Christopher Chiu, Micah T. McClain, Chris W. Woods, Geoffrey S. Ginsburg, Laura L. Elo, Ephraim L. Tsalik, Lara M. Mangravite, Solveig K. Sieberts

**Author notes:** These authors contributed equally (Slim Fourati, Aarthi Talla, Mehrad Mahmoudian, Joshua G. Burkhart, Riku Klén).

## Abstract

Respiratory viruses are highly infectious; however, the variation of individuals’ physiologic responses to viral exposure is poorly understood. Most studies examining molecular predictors of response focus on late stage predictors, typically near the time of peak symptoms. To determine whether pre- or early post-exposure factors could predict response, we conducted a community-based analysis to identify predictors of resilience or susceptibility to several respiratory viruses (H1N1, H3N2, Rhinovirus, and RSV) using peripheral blood gene expression profiles collected from healthy subjects prior to viral exposure, as well as up to 24 hours following exposure. This analysis revealed that it is possible to construct models predictive of symptoms using profiles even prior to viral exposure. Analysis of predictive gene features revealed little overlap among models; however, in aggregate, these genes were enriched for common pathways. Heme Metabolism, the most significantly enriched pathway, was associated with higher risk of developing symptoms following viral exposure.

---

Acute respiratory viral infections are among the most common reasons for outpatient clinical encounters *(1)*. Symptoms of viral infection may range from mild (e.g. sneezing, runny nose) to life-threatening (dehydration, seizures, death), though many individuals exposed to respiratory viruses remain entirely asymptomatic *(2)*. Variability in individuals’ responses to exposure has been observed both in natural infections *(3)* and controlled human viral exposure studies. Specifically, some individuals remained asymptomatic despite exposure to respiratory viruses, including human rhinovirus (HRV) *(4–6)*, respiratory syncytial virus (RSV) *(4–6)*, influenza H3N2 *(4–9)* and influenza H1N1 *(4, 5, 9)*. Factors responsible for mediating response to respiratory viral exposure are poorly understood. These individual responses are likely influenced by multiple processes, including the host genetics *(10)*, the basal state of the host upon exposure *(11)*, and the dynamics of host immune response in the early hours immediately following exposure and throughout the infection *(12)*. Many of these processes occur in the peripheral blood through activation and recruitment of circulating immune cells *(13)*. However, it remains unknown whether host factors conferring resilience or susceptibility to symptomatic infectious disease can be detected in peripheral blood before infection or whether they are only apparent in response to pathogen exposure.

In order to identify such gene expression markers of resilience and susceptibility to acute respiratory viral infection, we utilized gene expression data from seven human viral exposure experiments *(6, 7, 9)*. These exposure studies have shown that global gene expression patterns measured in peripheral blood around the time of symptom onset (as early as 36 hours after viral exposure) are highly correlated with symptomatic manifestations of illness *(6, 9)*. However, these later-stage observations do not necessarily reflect the spectrum of early time point immune processes that might predict eventual infection. Since transcriptomic signals are weak at these early time points, the detection of early predictors of viral response has not yet been possible in any individual study. By combining data collected across these seven studies and leveraging the community to implement state-of-the-art analytical algorithms, the *Respiratory Viral DREAM Challenge* (www.synapse.org/ViralChallenge) aimed to develop early predictors of resilience or susceptibility to symptomatic manifestation based on expression profiles that were collected prior to and at early time points following viral exposure.

## Results

### Human Viral Exposure Experiments

In order to determine whether viral susceptibility could be predicted prior to viral exposure, we collated 7 human viral exposure experiments: one RSV, two influenza H1N1, two influenza H3N2 and two human rhinovirus studies, in which a combined total of 148 healthy volunteers were exposed to virus (Fig. 1A-B) or sham (n=7) *(6, 7, 9)*. Subjects were excluded if pre-existing neutralizing antibodies were detected, except for the RSV study in which neutralizing antibodies were not an exclusion criteria. Each subject in the study was followed for up to 12 days after exposure and serially sampled for peripheral blood gene expression by Affymetrix Human U133A 2.0 GeneChips. Throughout the trial, subjects self-reported clinical symptom scores across 8-10 symptoms (Fig. S1); these data were used to stratify subjects as either symptomatic or asymptomatic and to quantify symptom severity. Additionally, nasopharyngeal swabs measured viral shedding; these data were used to stratify subjects as either shedders or non-shedders (Fig. 1C). Clinical symptoms were summarized based on a modified Jackson score *(14)* and viral shedding was determined to be present if two or more measurable titers or one elevated titer was observed within 24 hours following viral exposure *(15)*. Viral shedding and clinical symptoms were provided to *Respiratory Viral DREAM Challenge* teams only for the training data set (Fig. 1C). An additional, but not previously available, human exposure experiment to the RSV virus (n = 21) was used as an independent test data set (Fig. 1A). The study design for this data set was similar to those of the 7 original data sets.

**Fig. 1.**
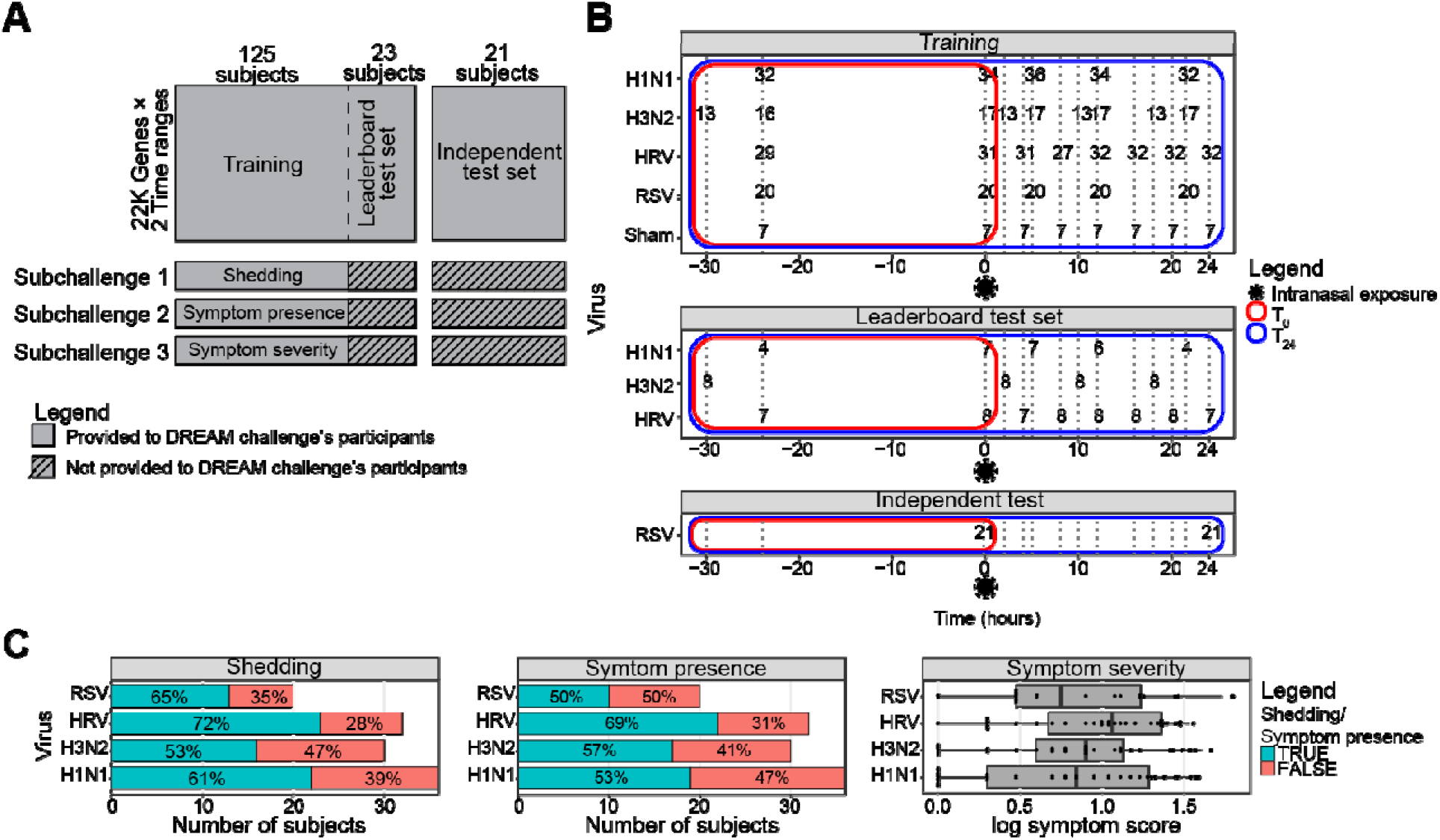
Respiratory Viral DREAM Challenge. **(A)** Schematic representation of the *Respiratory Viral DREAM Challenge*. **(B)** Challenge data come from seven viral exposure trials with sham or one of 4 different respiratory viruses (H1N1, H3N2, Rhinovirus, and RSV). In each of these trials, healthy volunteers were followed for seven to nine days following controlled nasal exposure to one respiratory virus. Blood was collected and gene expression of peripheral blood was performed 1 day (24 to 30 hours) prior to exposure, immediately prior to exposure and at regular intervals following exposure. Data were split into a training, leaderboard, and independent test set. Outcome data for the leaderboard and independent test set were not provided to the teams, but instead teams were asked to predict them based on gene-expression pre-exposure (T_0_) or up to 24 hours post-exposure (T_24_). **(C)** Symptom data and nasal lavage samples were collected from each subject on a repeated basis over the course of 7-9 days. Viral infection was quantified by measuring release of viral particles from viral culture or by qRT-PCR (“viral shedding”). Symptomatic data were collected through self-report on a repeated basis. Symptoms were quantified using a modified Jackson score, which assessed the severity of 8 upper respiratory symptoms (runny nose, cough, headache, malaise, myalgia, sneeze, sore throat and stuffy nose).

### Data Analysis Challenge

Using these data, an open data analysis challenge, the *Respiratory Viral DREAM Challenge*, was formulated. Teams were asked to predict viral shedding and clinical symptoms based on peripheral blood gene-expression from up to two timepoints: prior to viral exposure (T_0_) or up to 24 hours post viral exposure (T_24_). Based on gene expression data from the two timepoints, teams were asked to predict at least one of three outcomes: presence of viral shedding (subchallenge 1 (SC1)), presence of symptoms, defined as a modified Jackson score ≥ 6 (subchallenge 2 (SC2)), or symptom severity, defined as the logarithm of the modified Jackson score (subchallenge 3 (SC3)). Teams were asked to submit predictions based on gene-expression and basic demographic (age and gender) data from both timepoints to enable cross-timepoint comparison. The 7 collated data sets served as a training dataset on which teams could build their predictive models. For a subset of subjects (n = 23), phenotypic data were withheld to serve as a leaderboard test set for evaluation with real-time feedback to teams.

Teams were asked to submit at least one leaderboard submission at each timepoint to be evaluated on the leaderboard test set. Performance metrics for these models were returned in real-time, and teams could update their submissions accordingly up to a maximum of 6 combined submissions per subchallenge. At the end of this exercise, teams were asked to provide leave-one-out cross-validation-based predictions on the training set (LOOCVs) and predictor lists for each of their best models.

Submitted models were ultimately assessed on the held-out human RSV exposure data set that had not been publicly available, previously. Predictions for the binary outcomes (shedding and symptoms) were assessed using Area Under the Precision-Recall (AUPR) and Receiver Operating Characteristic (AUROC) curves, and ranked using the mean rank of these two measures. The predictions for the continuous outcome (symptom severity) were assessed using Pearson’s correlation with the observed values. In each case, permutation-based p-values were used to identify submissions that performed significantly better than those expected at random.

### Challenge Results

For presence of symptoms (SC2), 27 models were assessed; 13 models were developed using T_0_ predictors, and 14 models using T_24_ predictors. Four of the T_0_ models and three of the T_24_ models achieved a nominal p-value of 0.05 for AUPR or AUROC, with the best scoring models at each timepoint achieving similar scores (AUPR(T_0_)=0.958, AUROC(T_0_)=0.863, AUPR(T_24_)=0.953, AUROC(T_24_)=0.863). Team *Schrodinger’s Cat* was the only team that achieved significance for all measures and timepoints. Despite the few teams achieving statistical significance, the models submitted were overall more predictive than expected at random (enrichment p-values 0.008, 0.002, 0.021, and 0.05 for AUPR(T_0_), AUROC(T_0_), AUPR(T_24_), and AUROC(T_24_), respectively; Fig. 2A).

**Fig. 2.**
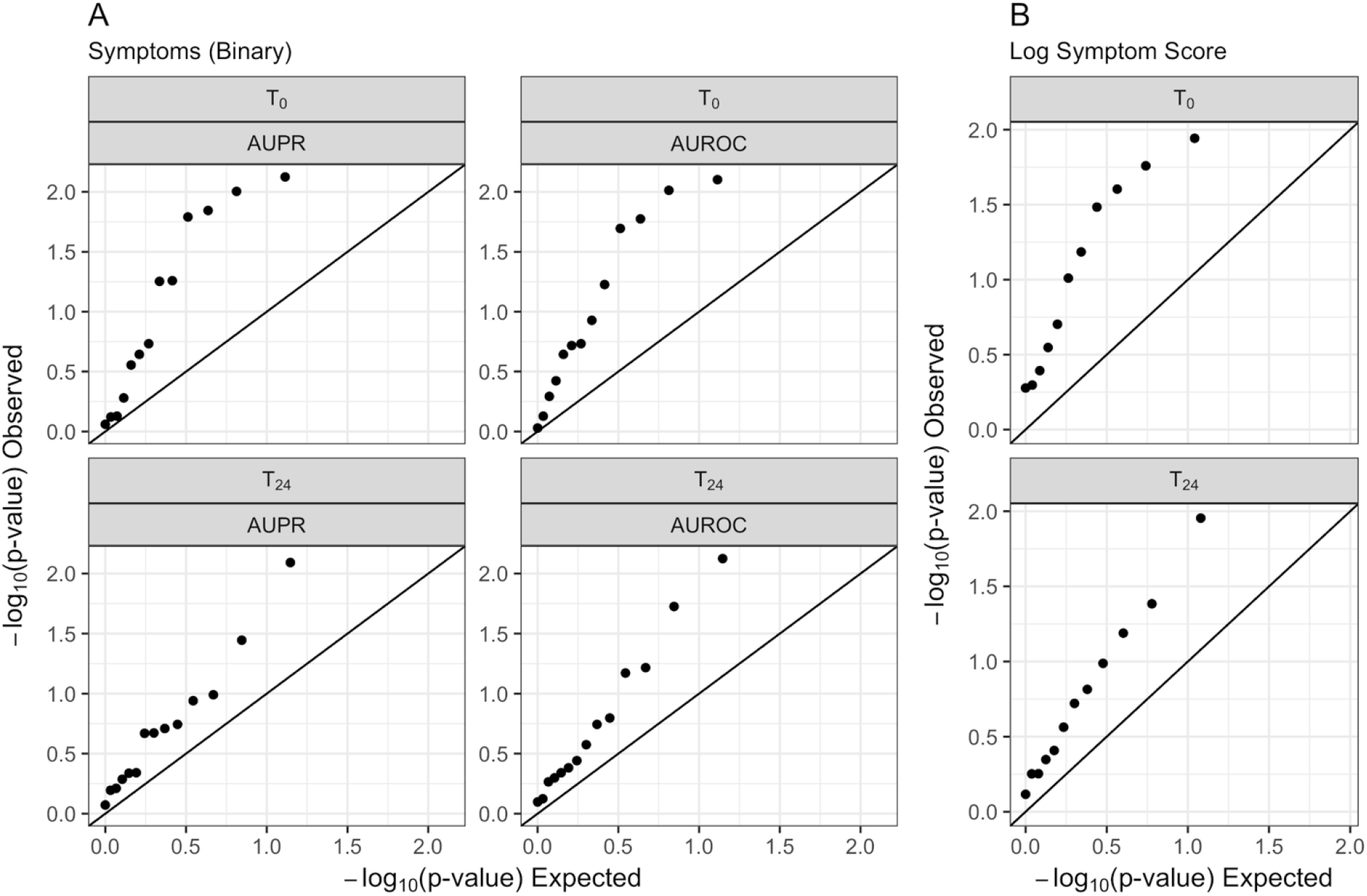
Models predict presence of symptoms and symptom severity better than expected at random. Observed −log_10_(p-value) versus the null expectation for submitted predictions for predicting **(A)** presence of symptoms (SC2) and **(B)** log symptom score (SC3). For both subchallenges significant enrichment of p-values (enrichment p-value 0.008, 0.002, 0.021, and 0.05 for AUPR(T_0_), AUROC(T_0_), AUPR(T_24_), and AUROC(T_24_), respectively, for presence of symptoms, and enrichment p-value 0.005 and 0.035 for T_0_ and T_24_, respectively, for log symptom score) across submissions demonstrates that pre- and early post-exposure transcriptomic data can predict susceptibility to respiratory viruses.

For symptom severity (SC3), 23 models were assessed; 11 models were developed using T_0_ predictors and 12 models using T_24_ predictors. Four of the T_0_ models and two of the T_24_ models achieved a nominal p-value of 0.05 for correlation with the observed log-symptom score, and as above, the best performing models scored similarly at both timepoints (r=0.490 and 0.495 for T_0_ and T_24_, respectively). Teams *cwruPatho* and *Schrodinger’s Cat* achieved significant scores at both timepoints. Consistent with SC2, we also saw that the models submitted were overall more predictive than expected at random (enrichment p-values 0.005 and 0.035 for T_0_ and T_24_, respectively; Fig. 2B). For both SC2 and SC3, enrichment was more pronounced at T_0_ compared to T_24_. Correlation between final scores and leaderboard scores was higher at T_0_, suggesting T_24_ predictions may have been subject to a greater degree of overfitting.

For viral shedding (SC1), 30 models were assessed from 16 different teams; 15 models were developed using T_0_ predictors and 15 models using T_24_ predictors. No submissions were statistically better than expected by random. In aggregate, these submissions showed no enrichment (enrichment p-values 0.94, 0.95, 0.82, and 0.95, for AUPR(T_0_), AUROC(T_0_), AUPR(T_24_), and AUROC(T_24_), respectively). In contrast, final scores were negatively correlated with leaderboard scores (correlation −0.22, −0.19, −0.65, and −0.54 for AUPR(T_0_), AUROC(T_0_), AUPR(T_24_), and AUROC(T_24_), respectively) suggesting strong overfitting to the training data or a lack of correspondence to viral shedding as assessed in the independent test data set, relative to the training data sets. The negative correlation was strongest at T_24_ (Fig. S2). Accordingly, results based on this subchallenge were excluded from further analysis.

### Best performing approaches

The two overall best performing teams were *Schrodinger’s Cat* and *cwruPatho*. Team Schrodinger’s Cat used the provided gene expression profiles before the viral exposure to predict shedding and log symptom scores (binary and continuous outcomes, respectively). For the T_0_ models, arithmetic means over measurements prior to exposure were calculated, whereas for the T_24_ models, only the latest measurements before viral exposure were used. Epsilon support vector regression (epsilon-SVR) *(16)* with radial kernel and 10-fold cross-validation were used to develop the predictive models. Their work demonstrated that predictive models of symptoms following viral exposure can be built using pre-exposure gene-expression.

Team *cwruPatho* constructed models of infection based on pathway modulation, rather than gene expression, to predict infection outcomes. To do so, they used a sample-level enrichment analysis *(17)* approach to summarize the expression of genes implicated in the Hallmark gene sets *(18)* of the Molecular Signature DataBase (MSigDB) *(19)*. They then fitted LASSO regularized regression models, which integrate feature selection with a regression fit *(20)*, on the pathways to predict shedding, presence of symptoms and symptom severity following viral exposure. Their work demonstrated that including multiple genes sharing the same biological function results in more robust prediction than using any single surrogate gene.

Teams *Schrodinger’s Cat* and *cwruPatho* used different feature transformation methods and machine learning approaches, suggesting that methods can successfully identify pre- or early post-exposure transcriptomic markers of viral infection susceptibility or resilience. To gauge the range of approaches taken, we extended this comparison to all *Respiratory Viral DREAM Challenge* teams who reported details on the methods they used to develop their submissions.

We assessed the range of data preprocessing, feature selection and predictive modeling approaches employed for the submissions, to determine whether any of these methods were associated with prediction accuracy. Details of these three analysis steps (preprocessing, feature selection and predictive modeling) were manually extracted from reports of 24 teams (35 separate reports) who submitted predictions either for the leaderboard test set or the independent test set. To more precisely reflect the conceptual variations across employed methodologies, each of these three analysis tasks was broken down into 4 data preprocessing categories, 7 feature selection categories and 9 predictive modeling categories (Table S1). Twenty of 24 (83.3%) teams employed some version of data preprocessing, the task most significantly associated with predictive ability (Fig. S3A). Specifically, exclusion of sham-exposed subjects and data normalization associated best with predictive performance (Fig. 3).

**Fig. 3.**
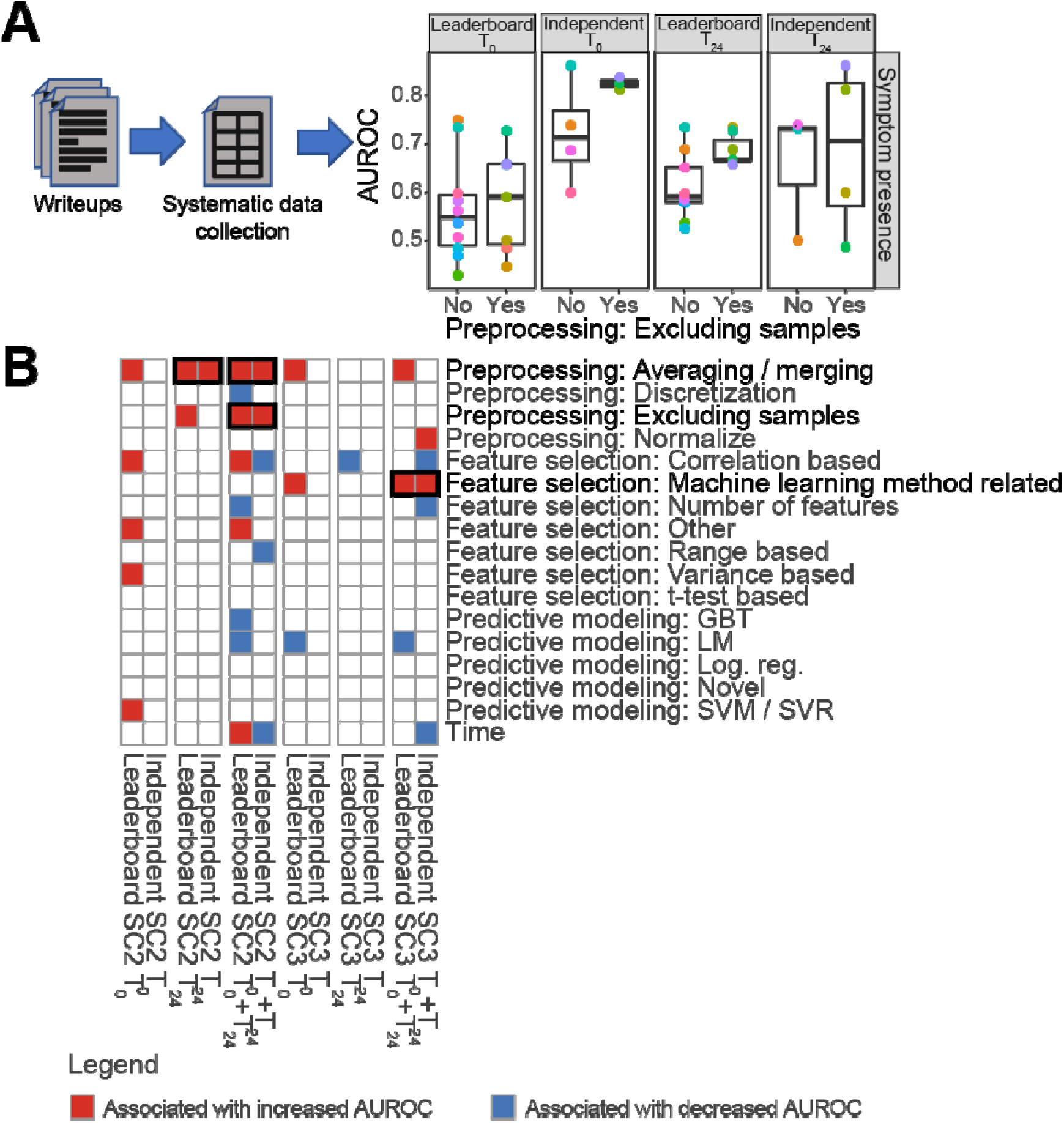
Adequate preprocessing leads to more accurate predictors of symptoms presence and severity. **(A)** Schematic representation of the analysis of the participating teams’ writeups to identify methodological steps associated with more accurate prediction of symptoms. First, the writeups were manually inspected to identify the preprocessing, feature selection and predictive modeling method used by each team. Second, the methods were regrouped into general categories across teams. Third, each general method was assessed for its association with predictive model accuracies on the leaderboard test set and the independent test set. **(B)** Heatmap showing the association of each general method with prediction ability (i.e. AUROC for subchallenge 2 (prediction of symptom presence; SC2) and Pearson’s correlation coefficient for subchallenge 3 (prediction of symptom severity; SC3)). For each general method, a Wilcoxon rank-sum test was used to assess the association between using the method (coded as a binary variable) and prediction ability.

Feature selection and predictive modeling approaches positively associated with predictive ability differed depending on whether the task was classification (presence of symptoms) or regression (symptom severity). Random forest-based predictive models performed slightly better than SVM/SVR methods at predicting symptom status (SC2) (Fig. S3B).

However, there was no discernible pattern relating feature selection and improved performance in SC2. Feature selection using machine learning approaches such as cross-validation was associated with improved performance in predicting symptom severity (SC3) (Fig. 3), as were SVM/SVR approaches when compared to linear regression model-based methods (e.g. logistic regression; Fig. S3C). Of note, SVM/SVR approaches were the most popular among the submissions.

We also sought to compare cross-timepoint predictions to determine the stability of predictions by timepoint. Significant correlation was observed between predictions using T_0_ and T_24_ gene expression for symptomatic classification (SC2) (Leaderboard: *ρ*=0.608, *p*=1.04e-61; Independent test set: *ρ*=0.451, *p*=2.05e-25). Interestingly, we observed that approximately 25% of subjects were difficult to predict based on T_0_ gene expression profile (inherently difficult; Fig. S4); similarly, approximately 25% of subjects were correctly predicted by the majority of teams (inherently easy; Fig. S4). Inherently difficult subjects were also misclassified when T_24_ gene expression data was used for the predictions. Inherently easy subjects were also consistently easy to classify using T_24_ gene expression data. This suggests *ab initio* characteristics allow some subjects to be more susceptible or resilient to symptomatic disease and that, within 24 hours, those characteristics are not substantially altered in post-exposure peripheral blood expression profiles.

### Biological Interpretation of Predictors

In addition to predictions, each team was asked to submit lists of gene expression features used in their predictive models. Six teams submitted separate models for each virus and reported virus-specific predictors. The remaining 28 teams reported predictors independent of virus, submitting a single model for all viruses. With the exception of the list from *cwruPatho*, which used pathway information in the selection of features, pathway analysis of individual predictor lists showed no enrichment of pathways from MSigDB *(19)*, possibly due to the tendency of most feature selection algorithms to choose one or few features from within correlated sets.

We then assessed whether models showing predictive ability (leaderboard test set AUROC > 0.5 for SC2 or r > 0 for SC3) tended to pick the same gene features, or whether the different gene sets may provide complementary information. Within each subchallenge and timepoint, significance of the overlap among predictor lists was calculated for every combination of two or more predictor lists across teams. All two-way, three-way, four-way, etc. overlaps were considered. This analysis revealed that there was no gene shared among all teams for any timepoint or subchallenge (Fig. 4A).

**Fig. 4.**
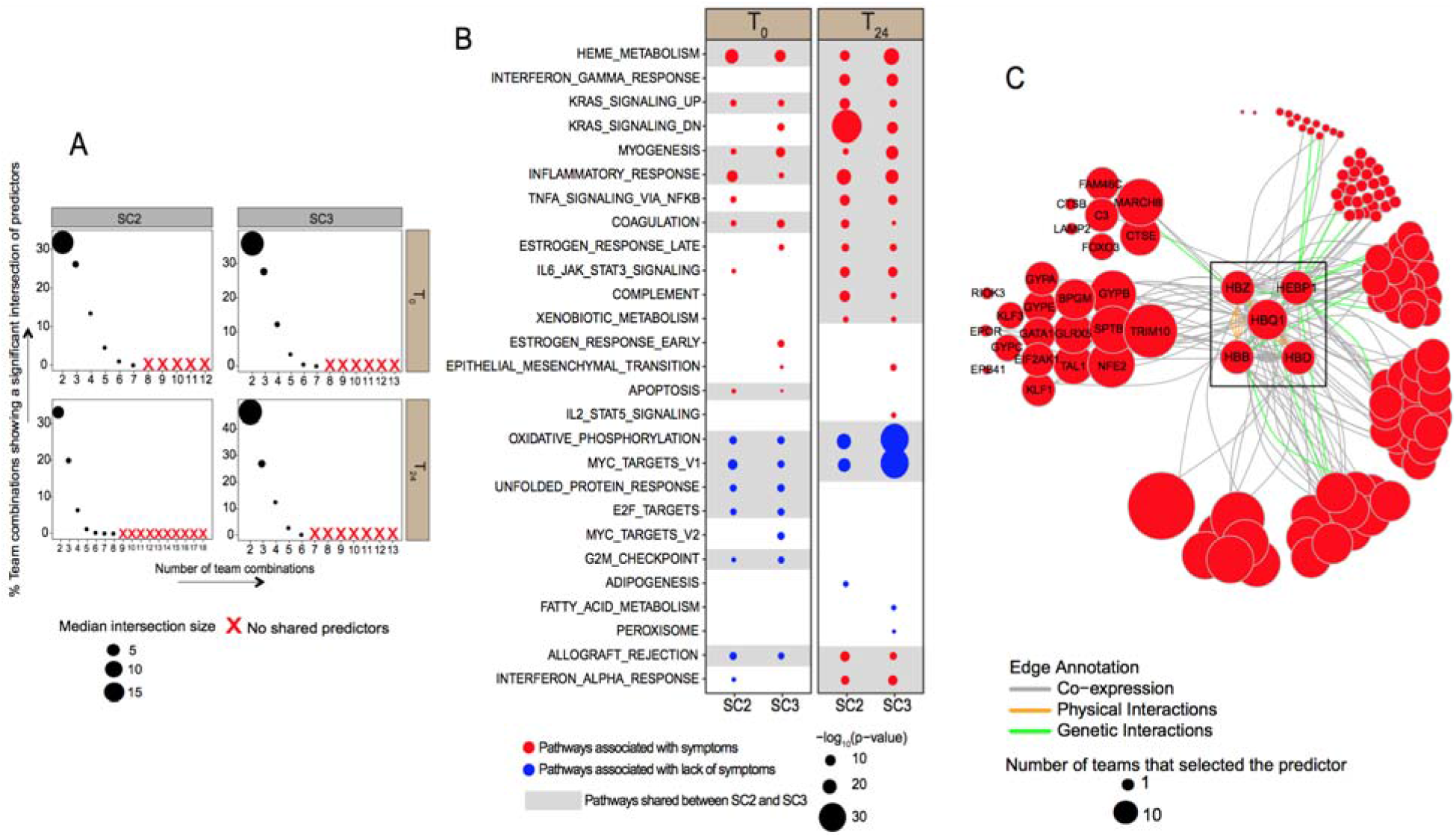
Overlap and pathway enrichment among predictors of symptoms. **(A)** Percent of team combinations showing statistically significant intersections of predictors at T_0_ and T_24_. Only teams whose with AUROC ≥ 0.5 or r ≥ 0 for subchallenge 2 and 3, respectively, were used for this analysis. The x-axis indicates the number of teams included in the combination. For example, the value 2 corresponds to pairwise overlaps, 3 corresponds to 3-way overlaps, etc. The y-axis indicates the percentage of team combinations with a statistically significant (p-value < 0.05) predictor intersection. Point size indicates median intersection size of predictors among team combinations with significant predictor intersection; ‘X’ indicates no significant predictor intersection. **(B)** Pathway enrichment among predictors of infection for each subchallenge (SC2 and SC3) at T_0_ and T_24_. The x-axis indicates subchallenge and each grid indicates timepoint. The y-axis indicates pathways enriched among predictors with a Benjamini-Hochberg corrected p-value < 0.05. Point size represents the fisher’s exact test enrichment −log_10_(p-value). Point colors indicate whether the pathway was associated with symptoms (red) or lack thereof (blue). Pathways shared between both SC2 and SC3 at each timepoint are highlighted in grey. Pathways are ordered by the decreasing maxP test statistic as determined in Fig S5 **(C)** GeneMANIA network of the union of predictors involved in the Heme metabolism pathway across time points (T_0_ and T_24_) and subchallenges (SC2 and SC3). Edges are inferred by GeneMANIA *(50)* corresponding to co-expression (purple), physical interactions (orange) and genetic interactions (green) among genes. Node size corresponds to the number of teams that selected the predictor.

Despite the paucity of overlap among predictor lists, we sought to identify whether genes used in the predictive models were part of the same biological processes or pathways. In other words, we examined if different teams might have chosen different surrogate genes to represent the same pathway. To test this hypothesis, we performed pathway enrichment analysis of the union of predictors across predictor lists within timepoint and subchallenge. We observed significant enrichments in each case (Fig. 4B), suggesting that predictive gene features are indeed complementary across models. More pathways were enriched among predictors from T_24_ models (SC2=17 pathways and SC3=20 pathways) than from T_0_ models (SC2=15 pathways and SC3=17 pathways). At T_0_, genes involved in the metabolism of heme and erythroblast differentiation (HEME METABOLISM), genes specifically up-regulated by KRAS activation (KRAS_SIGNALING_UP), genes defining an inflammatory response (INFLAMMATORY RESPONSE) and genes mediating cell death by activation of caspases (APOPTOSIS) were associated with presence of symptoms in both SC2 and SC3 (Fig. 4B). At T_24_, along with HEME METABOLISM, the expression of several inflammatory response pathways like KRAS SIGNALING, INFLAMMATORY RESPONSE, genes up-regulated in response to the gamma cytokine IFNg (INTERFERON GAMMA RESPONSE), genes upregulated by IL6 via STAT3 (IL6 JAK STAT3 SIGNALING), genes regulated by NFkB in response to TNF (TNFA SIGNALING VIA NFKB) and genes encoding components of the complement system (COMPLEMENT) were associated with symptoms in both SC2 and SC3 (Fig. 4B). Additionally, there was a significant overlap in genes across timepoints and subchallenges in each of these enriched pathways (Fisher’s exact test p-value ≤ 0.05) (Table S2).

A meta-analysis across subchallenges (SC2 and SC3) and timepoints (T_0_ and T_24_) was performed in order to identify the most significant pathways associated with outcome. HEME METABOLISM was the most significantly associated with developing symptoms (susceptibility), while OXIDATIVE PHOSPHORYLATION and MYC TARGETS were the most significantly associated with a lack of symptoms (resilience) (Fig. S5). This indicates that heme, known to generate inflammatory mediators through the activation of selective inflammatory pathways *(21)* is the best predictor of becoming symptomatic both pre- and early post-exposure to respiratory viruses. Genes in HEME METABOLISM associated with symptoms include genes coding for the hemoglobin subunits (HBB, HBD, HBQ1 and HBZ), the heme binding protein (HEBP1) and genes coding for enzymes important for the synthesis of heme (ALAS2, FECH, HMBS, UROD). It also includes glycophorins, which are the major erythrocyte membrane proteins (GYPA, GYPB, GYPC and GYPE), which are known receptors for the influenza virus (Fig. 4C) *(22, 23)*. Genes essential for erythroid maturation and differentiation (NEF2, TAL1, EPOR and GATA1), including the transcription factor GATA1 and its targets, the hemoglobin subunit genes HBB and HBG1/2, were also part of HEME METABOLISM associated with an increase in symptom frequencies and severity.

## Discussion

Using an open data analysis challenge framework, this study showed that models based on transcriptomic profiles, even prior to viral exposure, were predictive of infectious symptoms and symptom severity. The best scoring individual models for predicting symptoms and log-symptom score, though statistically significant, fall short of practical clinical significance. However, these outcomes suggest that there is potential to develop clinically relevant tests based on the knowledge gained from these results, though this would necessitate further efforts to generate more data or identify different biomarker assays which more accurately assess the mechanisms observed in the transcriptomic models.

A generally useful exercise in crowdsourcing-based challenges is to construct ensembles from the submissions to assimilate the knowledge contained in them, and boost the overall predictive power of the challenge *(24)*. This exercise has yielded useful results in earlier benchmark studies *(25, 26)* and the *DREAM Rheumatoid Arthritis Challenge (27)*. However, the ensembles constructed for the *Respiratory Viral DREAM Challenge* did not perform better than the respective best performers among all the individual submissions for the various subchallenges and time points. We attribute this shortcoming partly to the relatively small training set (118 subjects), which may incline the ensemble methods to overfit these data, and the assumption of class-conditioned independence of the submissions inherent in SUMMA may not have been appropriate in this challenge *(28)*. The relative homogeneity, or lack of diversity, among the submissions for the various subchallenges and timepoints may have been another potential factor behind the diminished performance of the ensembles *(29)*.

The relative homogeneity of submissions and observation that the same subjects are misclassified by almost all participating teams suggests there may be a plateau in predictive ability when using gene expression to predict the presence of symptoms or symptom severity. It is possible that an integrative analysis supplementing or replacing the gene expression data with post-transcriptional (such as metabolomic or proteomic) data could further improve accuracy.

For example, metabolomic data have been used to differentiate patients with influenza H1N1 from others with bacterial pneumonia or non-infectious conditions as well as differentiate influenza survivors from non-survivors *(30)*. With respect to proteomics, Burke *et al*. used four of the viral exposure studies described here to derive and validate a proteomic signature from nasal lavage samples which distinguish, with high accuracy, symptomatic from asymptomatic subjects at time of maximal symptoms *(31)*. Cytokines are a special class of proteins that has been investigated in a variety of infectious disease conditions. Of particular relevance, cytokine profiling has been performed for one of the influenza H3N2 studies used in this Challenge. In that work, McClain *et al*. demonstrated that several cytokines were upregulated early after viral exposure (within 24 hours in some cases) and differentiated symptomatic from asymptomatic cases *(32)*. Baseline differences in cytokine expression were not observed, however, suggesting that cytokine expression is useful for predicting response to viral exposure but not baseline susceptibility. To our knowledge, no study has identified baseline metabolomic or proteomic predictors of resilience or susceptibility to respiratory viral infection. In addition, the combination of these data with transcriptomic predictors has not yet been investigated and may yield robust predictors of susceptibility or resistance to infection.

Our analyses revealed a significant concordance between predictions at T_0_ and T_24_ (Fig. S4), as well as a significant overlap between predictors at each of these timepoints (Table S2). Given the stability of predictions and predictors between T_0_ and T_24_, it appears that the preexposure biological mechanisms conferring susceptibility or resilience to respiratory viral infection may be observable up to one day post-exposure. We also observed significant overlap between gene signatures at both T_0_ and T_24_ and late stage signatures of viral infection, reported in the literature, and derived from gene-expression 48 hours or later after viral exposure (Table S3) *(5–9, 15, 33–38)*. The overlap between the predictors identified in this study and the later stage signatures was more significant at T_24_ than T_0_, suggesting that pre-exposure signatures of susceptibility differ somewhat from post-exposure signatures of active infection, and T_24_ predictors may reflect some aspects of both. The T_0_ gene signatures may encompass novel insight into *ab initio* factors that confer resilience or susceptibility.

Pathway enrichment analysis in our study revealed that the most significantly enriched pathway associated with symptomatic infection was HEME METABOLISM, known to have a direct role in immunity through activation of innate immune receptors on macrophages and neutrophils *(21)*. Of note, genes part of HEME METABOLISM were also enriched among late stage signatures of viral infection (ex. Hemoglobin gene HBZ and the iron containing glycoprotein ACP5 in *(33))*. Iron (obtained from heme) homeostasis is an important aspect of human health and disease. Viruses require an iron-rich host to survive and grow, and iron accumulation in macrophages has been shown to favor replication and colonization of several viruses (e.g. HIV-1, HCV) and other pathogenic microorganisms *(39)*. Furthermore, iron-replete cells have been shown to be better hosts for viral proliferation *(39)*. Increased iron loading in macrophages positively correlates with mortality *(39)* and it has been shown that viral infection can cause iron overload which could further exacerbate disease. Additionally, previous evidence suggests counteracting iron accumulation may limit infection *(21, 39)*. Studies have shown that limiting iron availability to infected cells (by the use of iron chelators) curbed the growth of several infectious viruses and ameliorated disease *(21, 39–41)*. This important role of iron in the susceptibility and response to infection may be the mechanism by which HEME METABOLISM genes conferred susceptibility to respiratory viral infection. As such, it represents an important biological pathway potentially offering a means by which an individual’s susceptibility or response to infection can be optimized. Such a relationship should be investigated in future studies of infection susceptibility. In addition, Heme-oxygenase (HMOX1), a heme-degrading enzyme that antagonizes heme induced inflammation and is essential for the clearance of heme from circulation *(42)*, was among the predictors from the T_0_ models. Interestingly, the expression of this gene at baseline was associated with lack of symptoms (for both SC2 and SC3), in concordance with its reported antiviral role during influenza infection *(43, 44)*. Augmentation of HMOX1 expression by gene transfer had provided cellular resistance against heme toxicity *(45)*. Hence enhancing HMOX1 activity could be an alternative to antagonize heme induced effects and thereby controlling infection and inflammation.

In addition to HEME METABOLISM, pro-inflammatory pathways such as INFLAMMATORY RESPONSE, KRAS SIGNALING, and APOPTOSIS were also associated with susceptibility to viral infection in our study, while homeostatic pathways, such as OXIDATIVE PHOSPHORYLATION and MYC TARGETS, were associated with resilience, both prior to and post-viral exposure (Fig. 4). Enrichment of these pathways among T_24_ predictors was more significant than among the T_0_ predictors, suggesting these mechanisms are not only emblematic of baseline system health, but also response to viral invasion. Additional pathways enriched among T_24_ predictors include INTERFERON GAMMA RESPONSE and COMPLEMENT, which are involved in innate and acquired immunity. Several genes among T_0_ and T_24_ predictors overlapped with genes positively associated with flu vaccination response *(46)*. Among them, *FCER1G* and *STAB1*, members of the inflammatory response pathway positively associated with symptoms in this study and were elevated prior to vaccination in young adults who showed good response to vaccination *(46)* (Fisher exact test: p=0.0338 for T_0_ and p=0.000673 for T_24_). This suggest that individuals predicted at a higher risk of presenting symptoms following influenza exposure may also be the most likely to benefit from vaccination.

The *Respiratory Viral DREAM Challenge* is to date the largest and most comprehensive analysis of early stage prediction of viral susceptibility. The open data analysis challenge framework is useful for comparing approaches and identifying the most scientifically or clinically relevant model or method in an unbiased fashion *(24)*. In this case, we observed few commonalities among the best performing models of symptomatic susceptibility to respiratory viral exposure. Indeed, the overall best performing teams in the challenge used different machine learning techniques to build their models. Interestingly, data preprocessing was the analysis task most significantly associated with model accuracy, suggesting what has often been speculated, that adequate attention to data processing prior to predictive modeling is a crucial first step *(47)*.

The open data challenge framework is also useful in arriving at consensus regarding research outcomes that may guide future efforts within a field *(24)*. Through this challenge, we have identified *ab initio* transcriptomic signatures predictive of response to viral exposure, which has provided valuable insight into the biological mechanisms conferring susceptibility to infection. This insight was not evident from any individual model, but became apparent with the meta-analysis of the individual signatures. While development of a diagnostic test of baseline susceptibility in not yet feasible based on these findings, they suggest potential for development in this area.

## Methods

### Training Data

Training data came from seven related viral exposure trials, representing four different respiratory viruses. The datasets are *DEE1 RSV, DEE2 H3N2, DEE3 H1N1, DEE4X H1N1, DEE5 H3N2, Rhinovirus Duke*, and *Rhinovirus UVA (6, 7, 9)*. In each of these human viral exposure trials, healthy volunteers were followed for seven to nine days following controlled nasal exposure to the specified respiratory virus. Subjects enrolled into these viral exposure experiments had to meet several inclusion and exclusion criteria. Among them was an evaluation of pre-existing neutralizing antibodies to the viral strain. In the case of influenza H3N2 and influenza H1N1, all subjects were screened for such antibodies. Any subject with pre-existing antibodies to the viral strain was excluded. For the rhinovirus studies, subjects with a serum neutralizing antibody titer to RV39 > 1:4 at pre-screening were excluded. For the RSV study, subjects were pre-screened for neutralizing antibodies, although the presence of such antibodies was not an exclusion criterion.

Symptom data and nasal lavage samples were collected from each subject on a repeated basis over the course of 7-9 days. Viral infection was quantified by measuring release of viral particles from nasal passages (“viral shedding”), as assessed from nasal lavage samples via qualitative viral culture and/or quantitative influenza RT-PCR. Symptom data were collected through selfreport on a repeated basis. Symptoms were quantified using a modified Jackson score *(14)*, which assessed the severity of eight upper respiratory symptoms (runny nose, cough, headache, malaise, myalgia, sneeze, sore throat, and stuffy nose) rated 0-4, with 4 being most severe. Scores were integrated daily over 5-day windows.

Blood was collected and gene expression of peripheral blood was performed 1 day (24 to 30 hours) prior to exposure, immediately prior to exposure, and at regular intervals following exposure. These peripheral blood samples were gene expression profiled on the Affy Human Genome U133A 2.0 array.

All subjects exposed to influenza (H1N1 or H3N2) received oseltamivir 5 days post-exposure. However, 14 (of 21) subjects in the DEE5 H3N2 cohort received early treatment (24 hours postexposure) regardless of symptoms or shedding. *Rhinovirus Duke* additionally included 7 volunteers who were exposed to sham rather than active virus.

All subjects provided written consents, and each of the seven trials was reviewed and approved by the appropriate governing IRB.

### RSV Test Data

Healthy non-smoking adults aged 18-45 were eligible for inclusion after screening to exclude underlying immunodeficiencies. A total of 21 subjects (10 female) were inoculated with 10^4^ plaque-forming units of RSV A Memphis 37 (RSV M37) by intranasal drops and quarantined from 1 day before inoculation to the 12th day after. Peripheral blood samples were taken immediately before inoculation and regularly for the next 7 days and profiled on the Affy Human Genome U133A 2.0 array. Subjects were discharged after study day 12, provided no or mild respiratory symptoms and a negative RSV antigen respiratory secretions test. Shedding was determined by polymerase chain reaction (PCR) in nasal lavage and defined as detectable virus for ≥2 days between Day +2 and Day +10 to avoid false-positives from the viral inoculum and to align case definitions with the other 7 studies. Subjects filled a diary of upper respiratory tract symptoms from Day −1 to Day +12, which was summarized using a modified Jackson score. All subjects returned for further nasal and blood sampling on Day +28 for safety purposes. All subjects provided written informed consent and the study was approved by the UK National Research Ethics Service (London-Fulham Research Ethics Committee ref. 11/LO/1826).

### Analysis Challenge Design

The training data were split into training and leaderboard sets, where the leaderboard subjects were chosen randomly from 3 of the trials: *DEE4X H1N1, DEE5 H3N2*, and *Rhinovirus Duke*, which were not publicly available at the time of challenge launch. Outcome data for the leaderboard set were not provided to the teams, but instead, teams were able to test predictions in these individuals using the leaderboard, with a maximum of 6 submissions per subchallenge. Of these, at least one submission was required to use only data prior to viral exposure and at least one using data up to 24 hours post-exposure.

For the training data, teams had access to clinical and demographic variables: age, sex, whether the subject received early oseltamivir treatment (*DEE5 H3N2* only) and whether the subject received sham exposure rather than virus (*Rhinovirus Duke* only), as well as gene expression data for the entire time-course of the studies. They also received data for the three outcomes used in the data analysis challenge:

- Subchallenge 1: SHEDDING_SC1, a binary variable indicating presence of virus in nasal swab following exposure
- Subchallenge 2: SYMPTOMATIC_SC2, a binary variable indicating post-exposure maximum 5-day integrated symptom score >= 6
- Subchallenge 3: LOGSYMPTSCORE_SC3, a continuous variable indicating the log of the maximum 5-day integrated symptom score+1

as well as the granular symptom data by day and symptom category. For the leaderboard test data, they were supplied with the clinical and demographic variables and gene expression data up to 24 hours post-exposure.

Final assessment was performed in the RSV Test Data (*i.e*. independent test set), and outcomes for these subjects were withheld from teams. In order to assure that predictions were limited to data from the appropriate time window, the gene-expression data were released in two phases corresponding to data prior to viral exposure, and data up to 24 hours post exposure. Teams were also supplied with age and sex information for these subjects.

Both raw (CEL files) and normalized versions of the gene-expression data were made available to teams. Both versions contained only profiles that pass QC metrics including those for RNA Degradation, scale factors, percent genes present, β-actin 3’ to 5’ ratio and GAPDH 3’ to 5’ ratio in the *Affy* Bioconductor package. Normalization via RMA was performed on all expression data across all timepoints for the training and leaderboard data sets. The RSV data were later normalized together with the training and leaderboard data.

### Submission Scoring

Team predictions were compared to true values using AUPR and AUROC for subchallenges 1 and 2, and Pearson correlation for subchallenge 3. For each submission, a p-value, estimating the probability of observing the score under the null hypothesis that the predicted labels are random, was computed by 10,000 permutations of the predictions relative to the true values.

We also had access to leaderboard predictions from 10,000 models build on data with randomly permuted labels for 3 teams for SC2 and 2 teams for SC3. This second test estimates the probability of observing the score under the null hypothesis that the independent variables does not contain information about the target variable within the model structure used in the predictor. Comparisons between permutation p-values and scores from models built on the permuted data showed that the latter approach to p-value computation was slightly more conservative (data not shown), and presumably more robust to overfitting the training data. Albeit theoretically preferable, the computational demands of this approach makes it infeasible for most challenges.

### Heterogeneity of the Predictions

T_0_ and T_24_ predictions for each outcome and team were collected to assess whether they were correlated to each other. Three teams provided predictions as binary values while 12 teams provided predictions as continuous values on different scales. In order to compare binary and continuous predictions, we first transformed them into ranks (with ties given the same average rank) and then ordered subjects increasingly by their mean rank across outcomes (mean-rank). The lower the mean-rank, the more likely a subject was predicted by the teams as not showing shedding/symptoms, whereas a higher mean-rank means a subject was predicted by most of the teams as showing shedding/symptoms. Distribution of the mean-rank (Fig. S4) revealed three groups of subjects: (1) ~25% of subjects correctly predicted by most of the teams (i.e. inherently easy), (2) ~25% of subjects incorrectly predicted by most of the teams (i.e. inherently difficult) and (3) ~50% of subjects who were predicted differently by the teams.

### Ensemble Prediction

We constructed a variety of ensembles from the teams’ submissions to the various subchallenges as a part of the collaborative phase of the *Respiratory Viral DREAM Challenge*. To enable a comparative analysis between individual and ensemble models in the collaborative phase, the teams were requested to submit leave-one-out cross-validation (LOOCV)-derived predictions on the training examples using the same methods used to generate leaderboard and/or test set predictions in the competitive phase. The LOOCV setup, which doesn’t involve random subsetting of the training data, was chosen to avoid potential overfitting that can otherwise occur from training and testing on predictions made on the same set of examples *(25)*. We used three types of approaches for learning ensembles, namely stacking and its clustering-based variants *(25)*, Reinforcement Learning-based ensemble selection *(26)* methods, as well as SUMMA, an unsupervised method for the aggregation of predictions *(28)*. Consistent with the process followed by the individual teams, we learnt all the ensembles using the training set LOOCV-derived predictions described above, and used the leaderboard data to select the final models to be evaluated on the test data.

### Combined Gene Sets

Statistical significance of the overlap among predictor lists was calculated using the multi-set intersection probability method implemented in the SuperExactTest R package *(48)*. A first set of analysis was performed with teams whose leaderboard AUROC > 0.5. A second set of analysis aimed at identifying genes that overlap virus-specific, subchallenge-specific and timepoint-specific predictive models, was restricted to teams that provided virus-specific (*Nautilus, aydin, SSN_Dream_Team, Txsolo, cwruPatho* and *Aganita)*, subchallenge-specific (*aydin, SSN_Dream_Team, cwruPatho, jhou*) and timepoint-specific predictors (*aydin, SSN_Dream_Team, cwruPatho, Espoir, jdn, jhou, burkhajo*) and participated in the leaderboard phase of the challenge, respectively. For both analyses, overlapping predictors associated with p-values less than or equal to 0.05 were considered significant.

### Pathway enrichment analysis

To assess pathway enrichment among predictors of infection, we considered predictors from teams with leaderboard AUROC > 0.5 (SC2) or Pearson correlation, r > 0 (SC3). Affymetrix Human U133A 2.0 GeneChip probe identifiers were mapped to gene symbols. We removed probes matching multiple genes, and when multiple probes matched a single gene, we retained the probe with the maximum median intensity across subjects.

For the list of predictors of presence of symptoms (SC2), we calculated the log2 fold-change of features (symptomatic(1)/asymptomatic(0)) at T_0_ and T_24_, and for prediction of the symptom scores (SC3), we calculated the Spearman’s correlation coefficient of the features, at T_0_ and T_24_, with the outcome. Pathway enrichment was then performed on the union of all predictors (across the teams) that were associated with presence/increase severity of symptoms (SC2: log2 fold-change > 0 and SC3: Spearman’s correlation > 0), as well as, for the union of all predictors (across teams) that were associated with lack of symptoms/lower symptoms severity (SC2: log2 fold-change < 0 and SC3: Spearman’s correlation < 0), separately by timepoint and subchallenge. We used the Hallmark gene sets (version 6.0) *(18)* of the Molecular Signature DataBase (MSigDB) *(19)* for the enrichment, and calculated the significance of enrichment using Fisher’s exact test. The resulting p-values were corrected for multiple comparisons using the Benjamini and Hochberg algorithm. Only significantly enriched pathways (corrected p-value < 0.05) were reported. Meta-analyses across subchallenges and timepoints were performed using the maxP test statistic *(49)*.

## Acknowledgements

This work was supported by Defense Advanced Research Projects Agency and the Army Research Office through Grant W911NF-15-1-0107. The views expressed are those of the authors and do not reflect the official policy or position of the Department of Defense or the U.S.

Government. JGB was supported by a training grant from the National Institutes of Health, USA (NIH grant 4T15LM007088-25).GP and AS’s work was supported by NIH grant # R01GM114434 and an IBM faculty award to GP. TEMN was supported by the Ministry of Science and Technology of Taiwan grant 105-2218-E-006-016-MY2. KYY was supported by NIH grants U54 HL127624 and R01GM126019. MS was supported by Grants-in-Aid for Scientific Research JP16H02866 from the Japan Society for the Promotion of Science.

We wish to thank Rafick P. Sekaly (Case Western Reserve University) for his critical feedback during the writing process.

## Author Contributions

RH, CC, MTM, CWW, GSG, and ELT devised and performed the experiments. RH, GSG, ELT, LM and SKS designed and ran the data analysis challenge. SF, AT, MM, JGB, RK, RH, ZA, KYY, MEA, RA, SJ, XL, TEMN, MS, AS, RV, GP, LLE, SKS and The Respiratory Viral DREAM Challenge Consortium analyzed the data.

## Data Availability

Data are available through GEO GSE73072. Challenge results and methods and code for individual models are available through www.synapse.org/ViralChallenge.

## Materials & Correspondence

Correspondence should be addressed to Dr. Sieberts.

